# Proxi: a Python package for proximity network inference from metagenomic data

**DOI:** 10.1101/357764

**Authors:** Yasser EL-Manzalawy

## Abstract

Summary: Recent technological advances in high-throughput metagenomic sequencing have provided unique opportunities for studying the diversity and dynamics of microbial communities under different health or environmental conditions. Graph-based representation of metagenomic data is a promising direction not only for analyzing microbial interactions but also for a broad range of machine learning tasks including feature selection, classification, clustering, anomaly detection, and dimensionality reduction. We present Proxi, an open source Python package for learning different types of proximity graphs from metagenomic data. Currently, three types of proximity graphs are supported: *k*-nearest neighbor (*k*-NN) graphs; radius-nearest neighbor (*r*-NN) graphs; and perturbed *k*-nearest neighbor (*pk*-NN) graphs.

**Availability:** Proxi Python source code is freely available at https://bitbucket.org/idsrlab/proxi/.

**Supplementary information:** Tutorials and online documentation are available at https://proxi.readthedocs.io

## 1 Introduction

The study of microbial communities using high-throughput genomic surveys (e.g., 16S rRNA marker gene profiling) has been proven a powerful tool for probing the structure and diversity of microbial communities in different environments. Several computational methods based on statistical, machine learning, and network analysis approaches have been proposed. However, there is still a pressing need for more sophisticated methodologies and bioinformatics tools. Of particular interests are analysis tools for longitudinal and functional studies (Stulberg, et al., 2016).

An essential step for studying microbial interactions, which is the dominant driver of population structure and dynamics, is to identify correlations between taxa within ecological communities. Due to the compositional nature and the extreme sparsity of the metagenomic data, inferring correlation relationships is very challenging (Fang, et al., 2015; Friedman and Alm, 2012). Recently, a comparative study (Weiss, et al., 2016) showed that the state-of-the-art correlation detection tools have an extremely poor precision (i.e., below 0.2). Despite the existence of high rates of false positive edges in learned correlation networks, Abbas et al. (2018) showed that these networks could serve as a powerful tool for biomarkers discovery.

Motivated by this finding and by the widely acknowledged success of graph-based learning methods (Zhang, et al., 2013) in several application domains such as image processing, computer vision, and social network analysis, we present a Python tool for learning proximity graphs (Plaku and Kavraki, 2007) from metagenomic data. Proximity graphs are one of the most popular graph learning methods and are widely used in different machine learning tasks including (Zhang, et al., 2013): classification, clustering, dimensionality reduction, and anomaly detection. We believe that a tool for learning proximity graphs from metagenomic data is an essential tool for enabling the development of more sophisticated computational tools for biomarkers discovery, community detection, and longitudinal studies. We report a case study of identifying biomarkers for Inflammatory Bowel Diseases (IBD), using a benchmark dataset of 657 and 316 IBD and healthy controls metagenomic biopsy samples, respectively. Our results show that biomarker identification using the changes in nodes topological properties in proximity graphs outperforms some of the state-of-the-art feature selection methods.

## 2 Methods

Proxi constructs a proximity graph from the abundances of microbial operational taxonomic units (OTUs). Depending on the algorithm used and the user-specified parameters, the resulting graph could be weighted/unweighted and directed/undirected. Let *X ∈ R^p×n^* be an OTU table (in matrix representation) where each column corresponds a metagenomic sample and each row corresponds to an OTU. *x_ij_* represents the abundance (or relative abundance) of the *i^th^* OTU in the *j^th^* sample. Proxi learns a proximity graph *G = (V, E)* for *X* such that each node is an OTU and edges represent proximity relationships between nodes. The current implementation supports the construction of the following three nearest neighbor proximity graphs.

### 2.1 *k*-NN and *r*-NN graphs

Given an OTU table *X ∈ R^p×n^*, a *k*-NN graph *G = (V, E)* for *X* is a directed (or undirected) graph with |*V*| = *p* nodes such that there is an edge *e_ij_ ∈ E* if and only if *x_j_* is among the 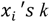 most similar OTUs using a distance metric *d(x_i_,x_j_*). Despite the simplicity of the procedure for constructing k-NN graphs, selecting an appropriate distance metric and deciding on the number of neighbors are not straightforward decisions. For small values of *k*, the resulting graph might not be connected (i.e., there might not be a path between every two vertices in the graph). Moreover, fixing the number of out-neighbors per node might not be an intuitive task. The *r*-NN graph is an alternative proximity graph where there is an edge *e_ij_ ∈ E* if and only if *d(x_i_,x_j_) ≤ r*. However, the choice of the value of *r* is data-dependent and it might be challenging to determine its appropriate value.

### 2.2 Perturbed *k*-NN graphs

The perturbed *k*-NN (*pk*-NN) graph construction algorithm (Wagaman, 2013) improves the construction of k-NN graphs from noisy data and yields proximity graphs that have a different *k* for each vertex. Briefly, the algorithm constructs *T* different *k*-NN graphs from the OTU table, *X*, by sampling with replacement from columns of *X*. A perturbed *k*-NN graph is then formed by aggregating the *T* graphs such that edges that appear at least in *cT* graphs are kept in the final graph, where 0 < *c* ≤ 1 is the graph aggregation tuning parameter.

### 2.3 Implementation and documentation

Proxi is implemented in Python 3.6 and requires NetworkX (Hagberg, et al., 2008) and Scikit-learn (Pedregosa, et al., 2011) to be installed. Constructed graphs could be saved in graphml or any NetworkX supported format. Similar to other graph inference tools (e.g., (Faust, et al., 2012; Kurtz, et al., 2015)), Proxi doesn’t support any network visualization functionality. However, any network analysis and visualization tool such as Cytoscape (Shannon, et al., 2003) could be directly used for visualization and downstream analysis of Proxi constructed graphs (for detailed examples, please see Proxi tutorials available at https://proxi.readthedocs.io/en/latest/Tutorials.html).

## 3 Results

Recently, Abbas et al. (2018) presented a novel Network-Based Biomarker Discovery (NBBD) framework for detecting disease biomarkers from metagenomic data. As a case study, we experimented with their NBDD framework and IBD train and test sets. The training set consists of 400 samples of equal number of IBD (positively labeled) and healthy (negatively labeled) samples. The test set consists of 457 and 116 IBD and healthy samples, respectively. Briefly, we applied the *pk*-NN graph construction algorithm using k=7, T=100, c=0.7, and Jaccard dissimilarity metric to infer (from training data) two *pk-*NN graphs corresponding to IBD and healthy populations, respectively. Then, a node importance score was computed using the absolute difference in node topological properties in the two graphs. Following Abbas et al. (2018), we considered the following node topological properties (all computed from the two *pk*-NN graphs using NetworkX library (Hagberg, et al., 2008)): Betweenness Centrality (btw); Closeness Centrality (cls); Average Neighbor Degree (and); Clustering Coefficient (cc); Node Clique Number (ncn); Core Number (cn). Fig. 1 compares the ROC curves estimated using the test data and Random Forests (RF) classifiers trained using different commonly used feature selection methods (top) and NBBD method (bottom). Interestingly, the network-based feature selection method using the change in node Closeness Centrality (cls) for scoring node importance yields a RF classifier with the highest AUC score of 0.81 whereas the top performing RF classifier using RF feature importance has an AUC score of 0.78.

**Fig. 1.**
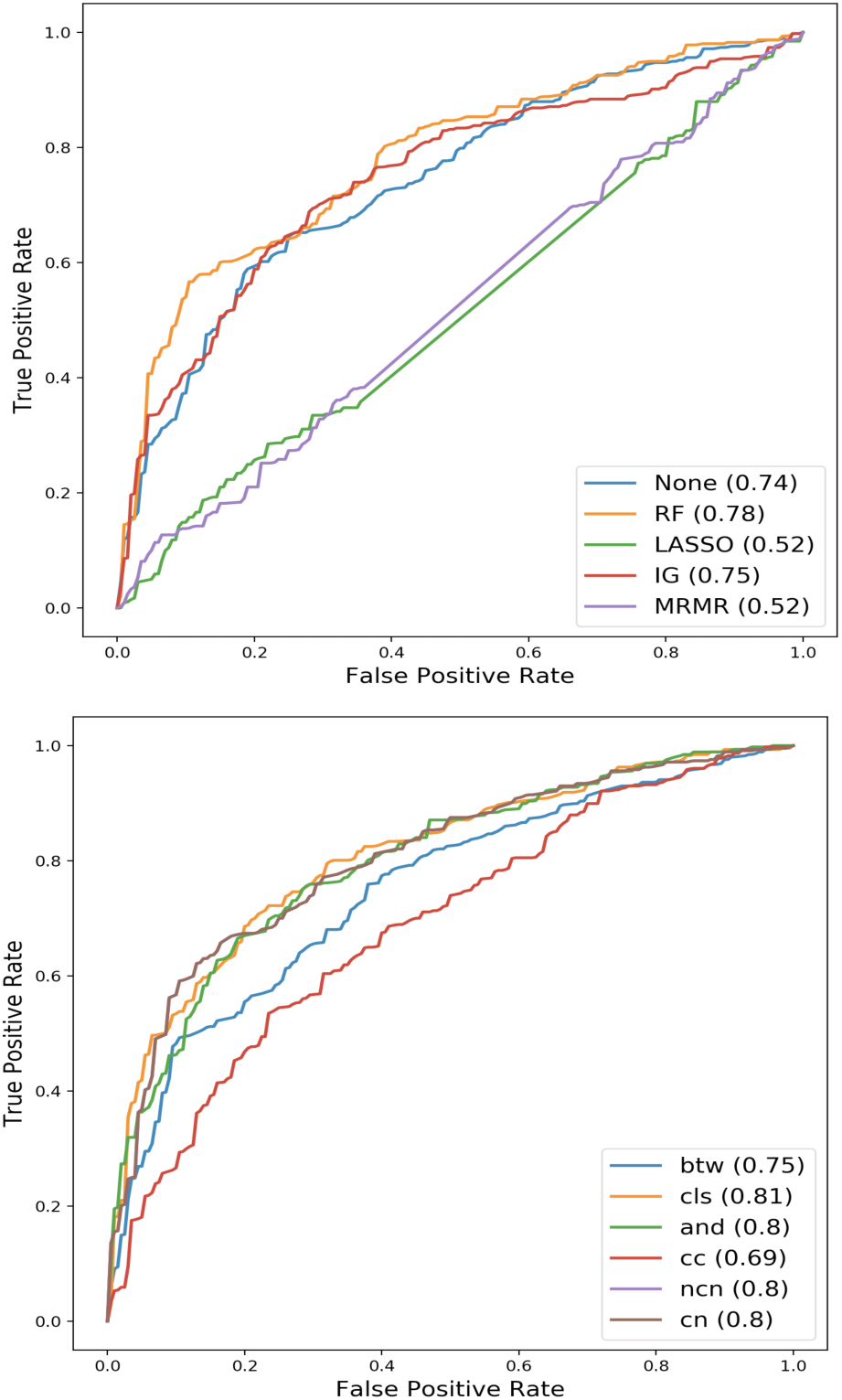
Performance comparison. ROC curves of RF classifiers evaluated using top 30 features selected using different machine learning based feature selection methods (top) and NBBD network-based feature selection method based on six node properties and pk-NN graphs. The AUC score for each classifier is shown between braces.

## 4 Conclusions

Application of machine learning algorithms to metagenomic data is challenging due to the extreme sparsity and the high-dimensionality of the metagenomic data. Learning a graphical representation of metagenomic data is a promising direction for modeling metagenomic data and enabling the application of a wide variety of well-developed graph mining algorithms for clustering, semi-supervised learning, and anomaly detection in metagenome-wide analysis studies.

We presented Proxi, an open source Python library for learning different types of proximity graphs from metagenomic data. The presented tool supports three types of proximity graphs: *k*-nearest neighbor (*k-*NN) graphs; radius-nearest neighbor (*r-*NN) graphs; and perturbed *k*-nearest neighbor (*pk-*NN) graphs. The *pk-*NN algorithm constructs improved *k-* NN graphs from noisy data using bootstrapping and graph aggregation techniques. We showed that the learned *pk-*NN graphs could be successfully used for identifying metagenomic biomarkers using the NBBD framework (Abbas, et al., 2018). In addition to our suggested method of using proximity graphs for biomarker identification (or graph-based feature selection), proximity graphs have been successfully used in literature for several machine learning tasks including: semi-supervised learning, clustering, anomaly detection, and manifold learning. Therefore, Proxi creates opportunities for researchers to explore and develop novel computational metagenomic analysis tools based on proximity graph models of the data.

## Acknowledgements

The author would like to thank Thanh Le, Ben Chen and Mostafa Abbas for testing Proxi and thoughtful discussions.

## References

Abbas, M., et al. Microbiomarkers Discovery in Inflammatory Bowel Diseases using Network-Based Feature Selection. In, Proceedings of the 9th ACM International Conference on Bioinformatics, Computational Biology, and Health Informatics. ACM; 2018. p. (in press). Preprint is available at: http://idsrlab.com/wp-content/uploads/2018/06/BCB18_YE.pdf

Fang, H., et al. CCLasso: correlation inference for compositional data through Lasso. Bioinformatics 2015;31(19):3172–3180.

Faust, K., et al. Microbial co-occurrence relationships in the human microbiome. PLoS computational biology 2012;8(7):e1002606.

Friedman, J. and Alm, E.J. Inferring correlation networks from genomic survey data. PLoS computational biology 2012;8(9):e1002687.

Hagberg, A., Swart, P. and S Chult, D. Exploring network structure, dynamics, and function using NetworkX. In.: Los Alamos National Lab.(LANL), Los Alamos, NM (United States); 2008.

Kurtz, Z.D., et al. Sparse and compositionally robust inference of microbial ecological networks. PLoS computational biology 2015;11(5):e1004226.

Pedregosa, F., et al. Scikit-learn: Machine learning in Python. Journal of machine learning research 2011;12(Oct):2825–2830.

Plaku, E. and Kavraki, L.E. Distributed computation of the knn graph for large high-dimensional point sets. Journal of parallel and distributed computing 2007;67(3):346–359.

Shannon, P., et al. Cytoscape: a software environment for integrated models of biomolecular interaction networks. Genome research 2003;13(11):2498–2504.

Stulberg, E., et al. An assessment of US microbiome research. Nature microbiology 2016;1(1):15015.

Wagaman, A. Efficient k-NN graph construction for graphs on variables. Statistical Analysis and Data Mining: The ASA Data Science Journal 2013;6(5):443–455.

Weiss, S., et al. Correlation detection strategies in microbial data sets vary widely in sensitivity and precision. The ISME journal 2016;10(7):1669.

Zhang, Y.-M., et al. Fast kNN graph construction with locality sensitive hashing. In, Joint European Conference on Machine Learning and Knowledge Discovery in Databases. Springer; 2013. p. 660–674.

